# Division-arrest induced filamentation protects uropathogenic *Escherichia coli* from killing by the cathelicidin antimicrobial peptide LL-37

**DOI:** 10.64898/2026.01.29.702582

**Authors:** Gozel Atakgayeva, Natasha Porco, Sofya Rudovskaya, Meghadeepa Sarkar, Madison Tomlinson, Raymond Huynh, Thomas J. Pucadyil, Roberto J. Botelho, Joseph B. McPhee

## Abstract

Antimicrobial peptides are a key component of the innate immune system, serving both as signaling and immunomodulatory molecules and also as direct antimicrobials toward numerous bacterial species. One of the key antimicrobial peptides in humans is LL-37 but the mechanism of bacterial killing remains poorly characterized. Using live-cell imaging, we observe that LL-37 preferentially targets actively dividing *E. coli* cells. Permeabilization occurs during late-stage division at the bacterial septum with both of the newly dividing cells being killed at the same time. Death occurs downstream of the recruitment and dissembly of FtsZ and FtsN and concomitantly with AmiB association at the divisome, consistent with killing occurring during late division. Model membrane tubes that topologically mimic the late-stage cytoplasmic membrane intermediates during division also undergo rapid size- and lipid composition-dependent fission, which likely reflect the ability of LL-37 to disrupt membranes. Intriguingly, we show that inducing bacterial filamentation results in resistance to LL-37 mediated killing. Resistance to LL-37 appears to be a general feature of bacterial filamentation as overexpression of *queE*, *sulA* or *damX* all resulted in reduced killing by LL-37. As filamentation is often observed during UPEC infection, we hypothesize that filamentation associated resistance to host innate immunity may be a key feature of UPEC pathogenesis that protects bacteria during extrusion from infection bladder epithelial cells, when exposure to soluble and cellular host defences is likely highest. This observation may provide an important avenue for therapeutic intervention.

**Significance:** Antimicrobial peptides are a key component of innate immunity. The human cathelicidin peptide, LL-37, is membrane active but the mechanism of sensitization in living systems is poorly understood. We show that dividing *E. coli* cells are highly susceptible to LL-37 mediated disruption. Killing occurs downstream of the assembly and activation of functional divisomes. In vitro membrane nanotubes also demonstrate sensitivity to LL-37 in a lipid composition and diameter-dependent manner. Cells with stalled division are protected from LL-37. Urinary tract infections caused by uropathogenic *E. coli* (UPEC) are a major cause of morbidity globally. UPEC form filamentous bacteria late in the human infection cycle. UPEC filaments are protected from LL-37 killing and suggest that interfering with bacterial filamentation might potentiate innate immune control of UTIs.

## Introduction

Antimicrobial peptides (AMPs) are found throughout all domains of life, serving to protect the producing organism from competition with or infection by susceptible microbes (1). In humans, AMPs are essential components of the innate immune system, providing a robust and rapid first line of protection from invading pathogens. Typically small (<50 amino acids) and cationic, they are produced by phagocytes like neutrophils and macrophages as well as by epithelial cells. In addition to their directly antimicrobial activity, AMPs can also modulate immune responses by recruiting immune cells, promoting wound healing and by regulating inflammatory responses in the host (2). Humans produce a number of different types of antimicrobial peptides, including α- and β-defensins, RegIII family AMPs, histatins and the only member of the cathelicidin family of AMPs, LL-37 (3).

LL-37 is synthesized as a pre-propeptide (hCAP-18) that is targeted for secretion before being cleaved by host proteases to the active LL-37 peptide. LL-37 is unstructured in aqueous solutions but takes on a curved α-helical structure in membrane-mimetic solvents (4). LL-37 exhibits antimicrobial activity against a number of human pathogens including, *Staphylococcus aureus*, *Enterococcus spp*., *Listeria monocytogenes*, *Escherichia coli*, *Salmonella enterica* and *Klebsiella pneumoniae* (5). The antimicrobial activity of LL-37 and that of many other AMPs depends on the ability of that peptide to interact with and disrupt target membranes (6–8). Despite advances in understanding the mechanisms by which LL-37 interacts with bacterial membranes and disrupts them leading to cell death, there remain significant gaps in our understanding of whether specific membrane components are targeted and precisely how LL-37 mediated bacterial death occurs.

Here, we show that dividing *E. coli* are selectively killed by LL-37. Using single-cell killing assays, we demonstrate that LL-37-mediated bacterial death is associated with late bacterial division concomitant with recruitment of late divisome components and after membrane invagination occurs. Altering the site of bacterial division also alters the point at which LL-37 mediated membrane disruption occurs. Furthermore, nanotube models of membrane structures are significantly more sensitive to LL-37-mediated cleavage as they thin. LL-37-mediated nanotube fission is also altered by lipid composition. We also demonstrate that genetic induction of filamentation by overexpression of *queE*, *sulA* or *damX* all result in enhanced LL-37 resistance in uropathogenic *E. coli*, suggesting that protection from killing is a conserved feature of non-dividing *E. coli*. This study provides substantial support for a model whereby bacterial division stalling during UPEC infection may be a bet-hedging strategy serving to protect a subpopulation of bacteria from the host-derived cellular and soluble immune factors and provides a rationale for developing strategies to target this response.

## Results

### Actively dividing cells are selectively killed by LL-37

Many previous results have established that killing of bacterial cells by LL-37 involves disruption of membrane integrity, but the precise mechanism of killing in living cells remains enigmatic. Actively dividing mid-logarithmic phase *E. coli* BW25113 cells were washed before being placed on lysogeny broth agarose pads in the presence of 1 µg/ml propidum iodide or 1 μg/ml SYTOX Green. LL-37 dissolved in water was then added to the edge of the cover slip and cells were monitored for up to one hour. We noted substantial preferential killing of bacteria that were actively dividing at the time of LL-37 exposure (Fig. 1). Furthermore, both SYTOX Green and PI dye entry was preferentially associated with the site of bacterial division, appearing to enter at the division site before rapidly diffusing throughout the cytosol of both newly divided daughter cells (Fig. 1 and Supplemental Movie S1). We also noted that division-associated killing was not unique to *E. coli* as we observed similar results with dividing *Salmonella enterica* (Supplemental Fig. S1)

**Figure 1.**
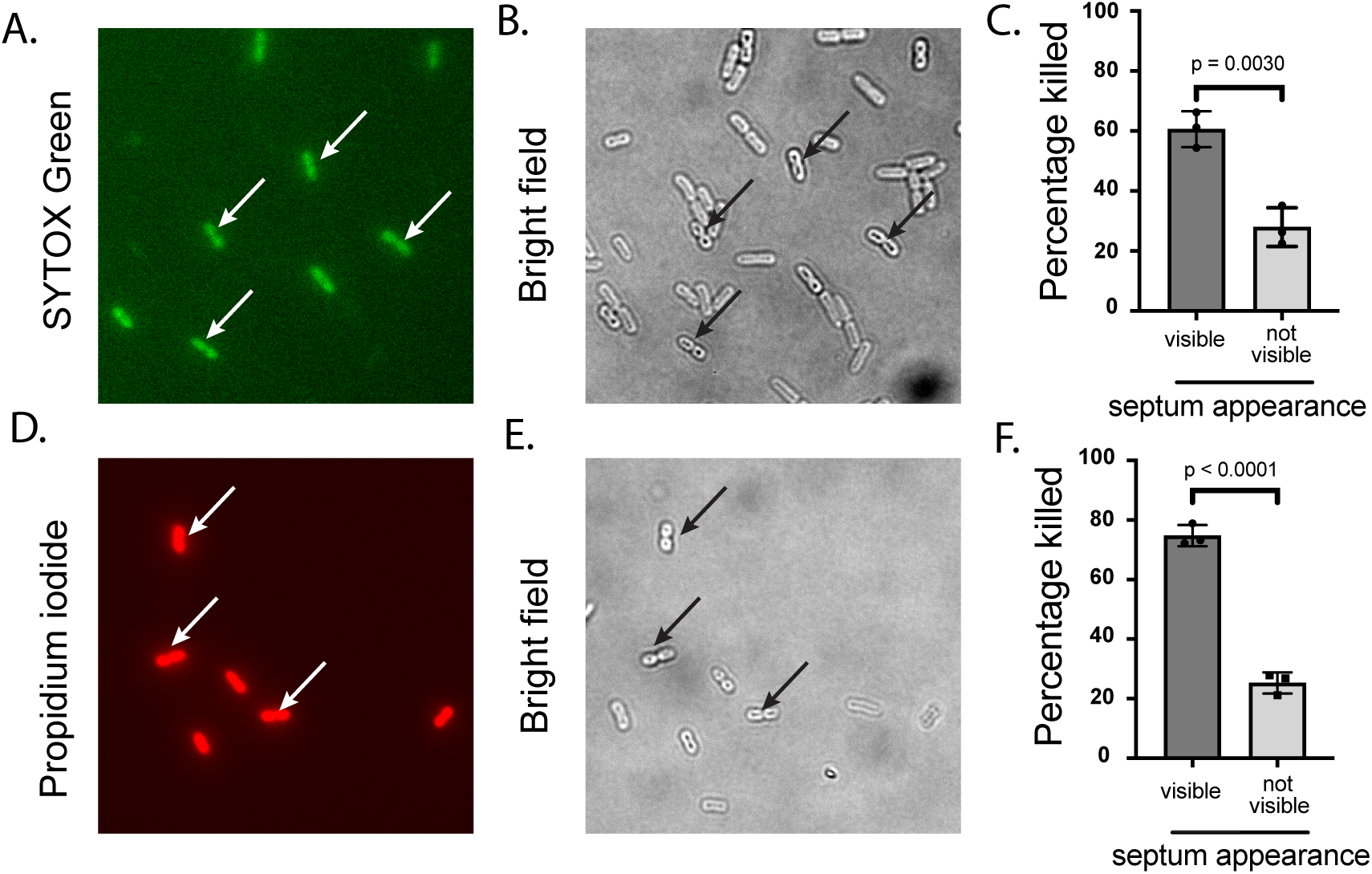
LL-37 preferentially permeabilizes actively dividing cells. *E. coli* BW25113 cells were grown in M9-glucose minimal medium and mid-logarithmic phase bacteria were then placed on 1% agarose pads with A-C) SYTOX™ Green nucleic acid stain or D-E) propidium iodide. Live images were collected while 3 µl of 50-100 µg/ml LL-37 in water was added to the edge of the agarose pad. Arrows indicate dead cells with evidence of a mid-cell invagination consistent with the formation of a divisome A&B; C&D. Statistical significance was tested by unpaired Student’s t-test comparing the average percentage of dead cells that contained a visible mid-cell invagination consistent with divisome formation and activation from three independent biological replicates. N = 141 to 308 dead cells per replicate measured via SYTOX Green staining and N = 77 to 232 dead cells per replicate for PI mediated killing.

### Division associated LL-37 killing occurs following recruitment, assembly and dissassembly of FtsZ and FtsN from the division site

To better understand the mechanistic basis of division-associated LL-37 killing, we created FtsZ-GFP, FtsN-SPOR-GFP and AmiB-sfGFP fusions and observed the kinetics of LL-37-mediated killing in relation to divisome-associated GFP/sfGFP recruitment and disassembly. Consistent with previous results, we observed assembly of FtsZ and FtsN-SPOR rings at the midline of actively dividing bacteria. In the presence of LL-37, cells that are actively dividing take up propidium iodide immediately following the disassembly of the mid-cell FtsZ and FtsN rings Fig. 2A-D. We observe that a larger proportion of AmiB-sfGFP appears to remain at the midline at the time of PI uptake but in general, killing follows disappearance of divisome-associated markers (Fig. 2E-F). Non-dividing cells in the observed fields of view were not killed at appreciable rates under these conditions (Supplemental Movies S1-S3).

**Figure 2.**
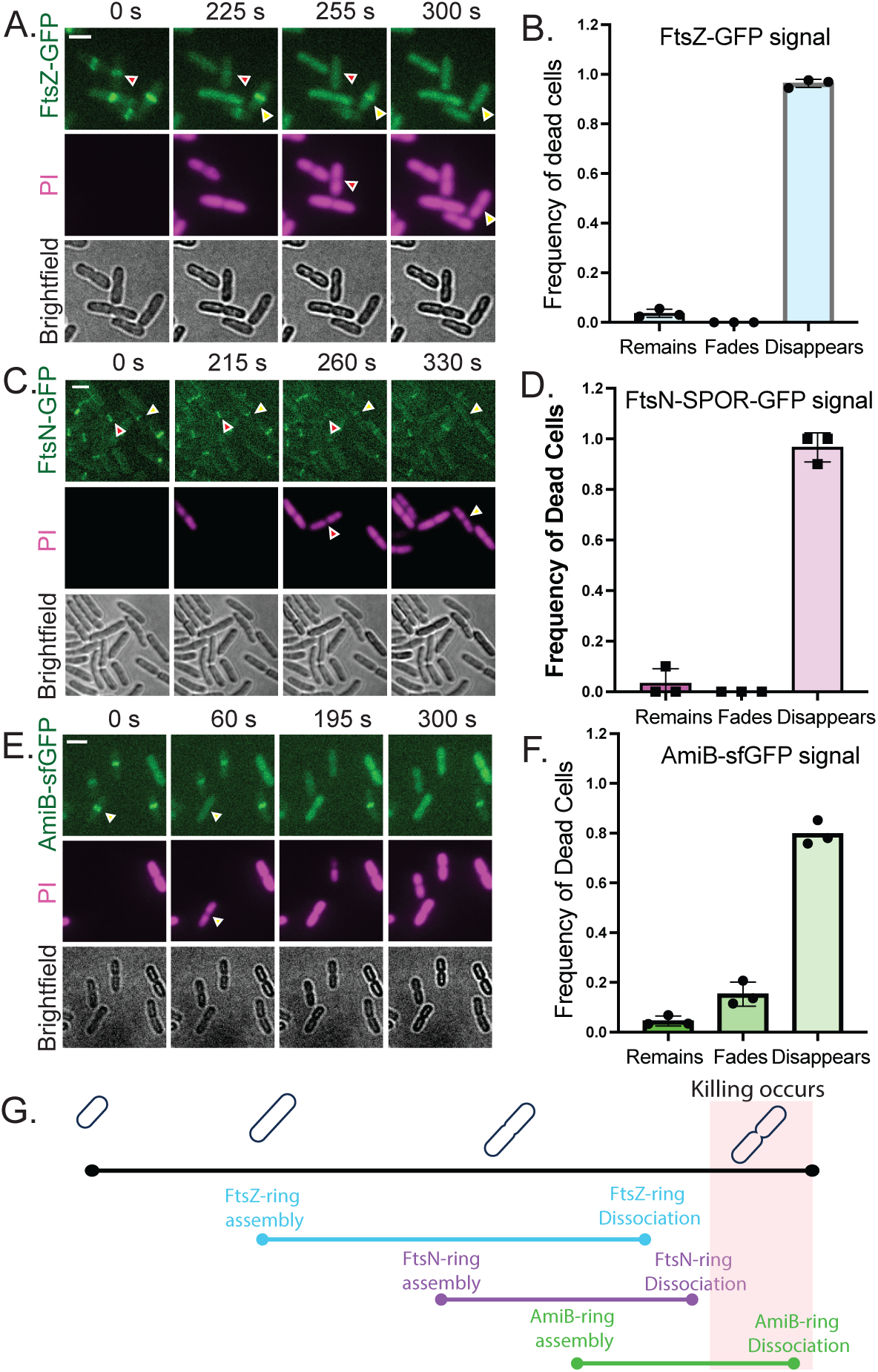
- LL-37 killing occurs downstream of FtsZ and FtsN recruitment and disassembly and concomitantly with septal closure and AmiB recruitment. A) Imaging of propidium iodide uptake in BW25113 cells expressing FtsZ-GFP exposed to LL-37 B) Characterization of FtsZ-GFP behaviour in LL-37 killed cells C) Imaging of propidium iodide uptake in BW25113 cells expressing FtsN-SPOR-GFP exposed to LL-37 D) Characterization of FtsN-SPOR-GFP behaviour in LL-37 killed cells E) Imaging of propidium iodide uptake in BW25113 cells expressing AmiB-sfGFP exposed to LL-37 F) Characterization of AmiB-sfGFP behaviour in LL-37 killed cells. To assess dying cells (B, D, E), PI+ cells were categorized based on the behaviour of the divisome-localized marker at the time of LL-37 mediated killing. Three biological replicates were assessed in each experiment. N = 29 to 132 dead cells counted for each biological replicate. G) schematic relating the timing of LL-37-mediated killing to the assembly and dissassembly of selected divisome protein marker behaviour.

### Mislocalization of division site alters location of LL-37 mediated cell death

As we observed a correlation between the maturation of the divisome and midcell-associated bacterial killing, we wanted to further test whether this was due to the location of the divisome or some other mid-cell associated property. In *E. coli*, the site of divisome initiation and assembly is controlled by the MinCDE system. MinC is an inhibitor of FtsZ polymerization. The MinDE proteins oscillate along the long axis of the cells, recruited to polar regions via a charged membrane targeting sequence that preferentially associates with polarly enriched cardiolipin and/or other anionic lipid species (9, 10). Polar localization of MinC inhibits assembly of FtsZ near the poles, thereby promoting midline association of FtsZ. Overexpression of FtsZ or deletion of *minCD* leads to increased rates of aberrant cell division and the formation of minicells due to initiation and activation of polarly-localized divisomes (11, 12). Consistent with previous reports, deletion of *minCD* or overexpression of *ftsZ* results in altered size distribution of *E. coli* cells, with increased numbers of minicells as well as a larger proportion of cells that are elongated (Fig. 3A). The shift to filamentation is more pronounced in the *ftsZ* overexpressing strain while cells lacking *minCD* show enhanced production of both filaments and minicells, with the Δ*minCD* cells showing a significantly greater proportion of cells smaller than 0.5 <u>μm in length (WT - 0%, Δ*minCD* - 19.3%, WT + pBAD*ftsZ* - 0.45%)</u>. We monitored LL-37 mediated killing in live *E. coli* K12 or K12Δ*minCD* cells and noted that LL-37 mediated killing occurred at both midline and minicell/polar divisomes with propidium iodide uptake observed from both locations (Fig 3B). Whereas WT *E. coli* showed almost exclusive killing via mid-cell associated divisomes, the Δ*minCD* strains showed permeabilization/killing from both polarly localized divisomes as well as from mid-cell associated divisomes (Fig 3B, 3D).

**Figure 3.**
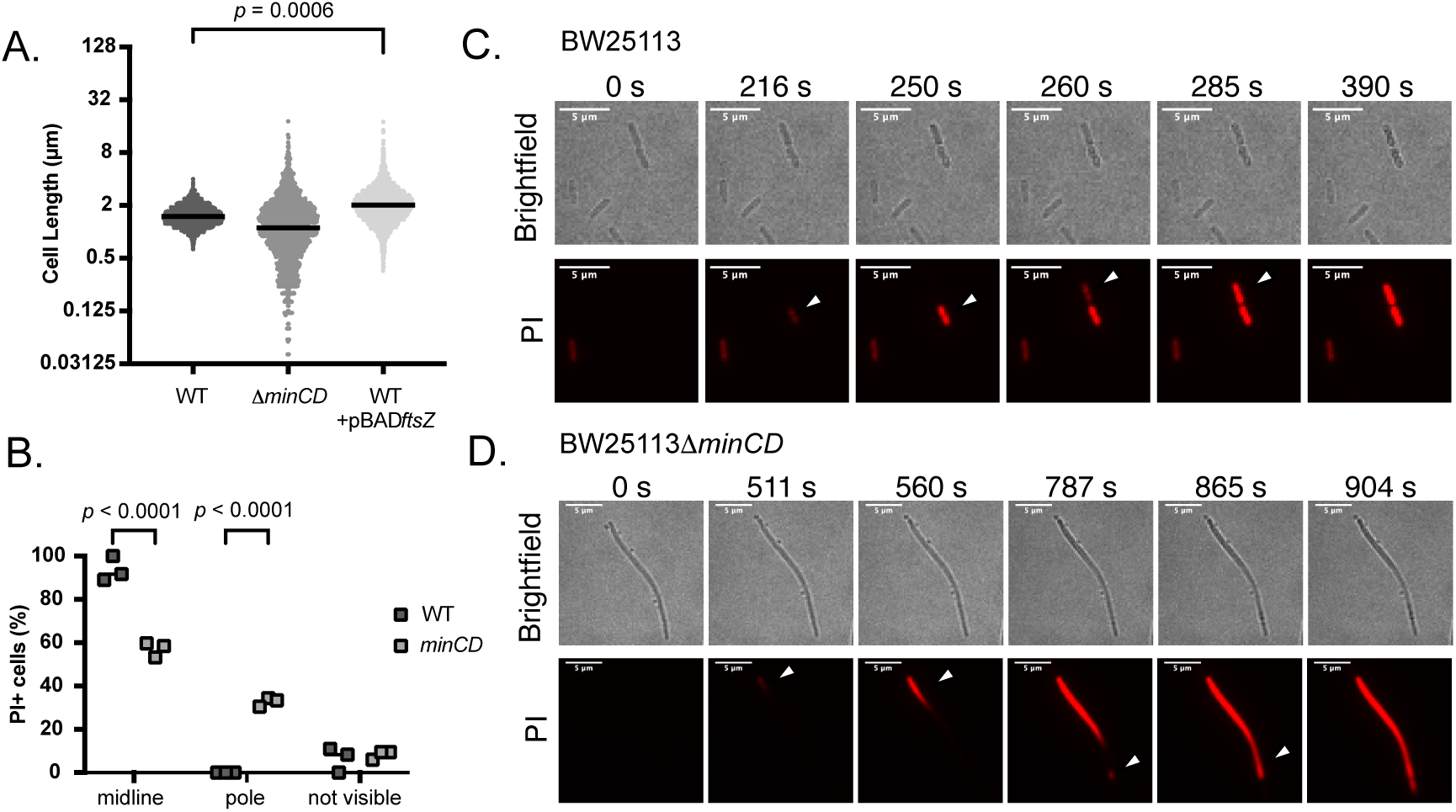
Induction of minicells via deletion of *minCD* leads to aberrant polar localization of the divisome site and induces LL-37-mediated killing from non-septal regions. A) Deletion of *minCD* or overexpression of FtsZ alters *E. coli* cell length and the distribution of cell lengths within the population. At least 1700 cells total were measured per strain and cells from three independent experiments were measured. The means of lengths measured in each experiment were compared by a log normal Brown-Forsynthe and Welch’s ANOVA with Dunnett’s T3 post-hoc testing. B) BW2511311*minCD* cells show LL-37 mediated killing from polar division sites. The site of LL-37-mediated killing of *E. coli* K12 vs K1211*minCD* bacteria were compared via two-way ANOVA with Šídák’s post-hoc test to determine whether different strains showed different sites of killing C) dying BW25113 or D) filamentous *E. coli* BW2511311*minC*D cell upon exposure to LL-37 in the presence of propidium iodide

### LL-37 leads to increased lipid nanotube fission in a diameter-dependent manner

As the majority of evidence to date suggests that LL-37 works via a membrane specific interaction, we considered the role of membrane morphology in sensitivity to disruption by LL-37. During bacterial cell division, a number of changes occur to the cytoplasmic membrane including that the relatively planar cytoplasmic membrane, with a large radius of curvature, adopts a narrowing tubular morphology, with a shrinking radius of curvature, at the septum during division-associated septal thinning. To mimic the thin tubular membrane topologies seen during division, we used supported membrane templates (SMrTs). (13). Templates were prepared with an *E. coli* polar lipid extract (PLE) doped with trace amounts (1 mol%) of Texas Red DHPE for visualization. Flowing 50 nM LL-37 into the system showed no apparent effect on the planar bilayer (Supplemental Fig. S2A), but tubular membranes underwent rapid and catastrophic severing (Fig. 4A, Supplemental Movie S4). Time lapse imaging showed that LL-37 induced alternating regions of dim and bright fluorescence along the length of the tube before fission occurs (Fig 4C). Membrane tubes in these preparations were estimated to be ∼12 ± 3 nm radius (mean ± SD, n = 11, Fig. 4A). Indeed, LL-37 caused a tube with a starting radius of ∼10 nm to undergo localized thinning, down to ∼3 nm (Fig. 4C, red arrowhead, Fig. 4D red trace) with the adjacent region concomitantly bulging to ∼25 nm (Fig. 4C, black arrowhead, Fig. 4D black trace) before fission.

**Figure 4.**
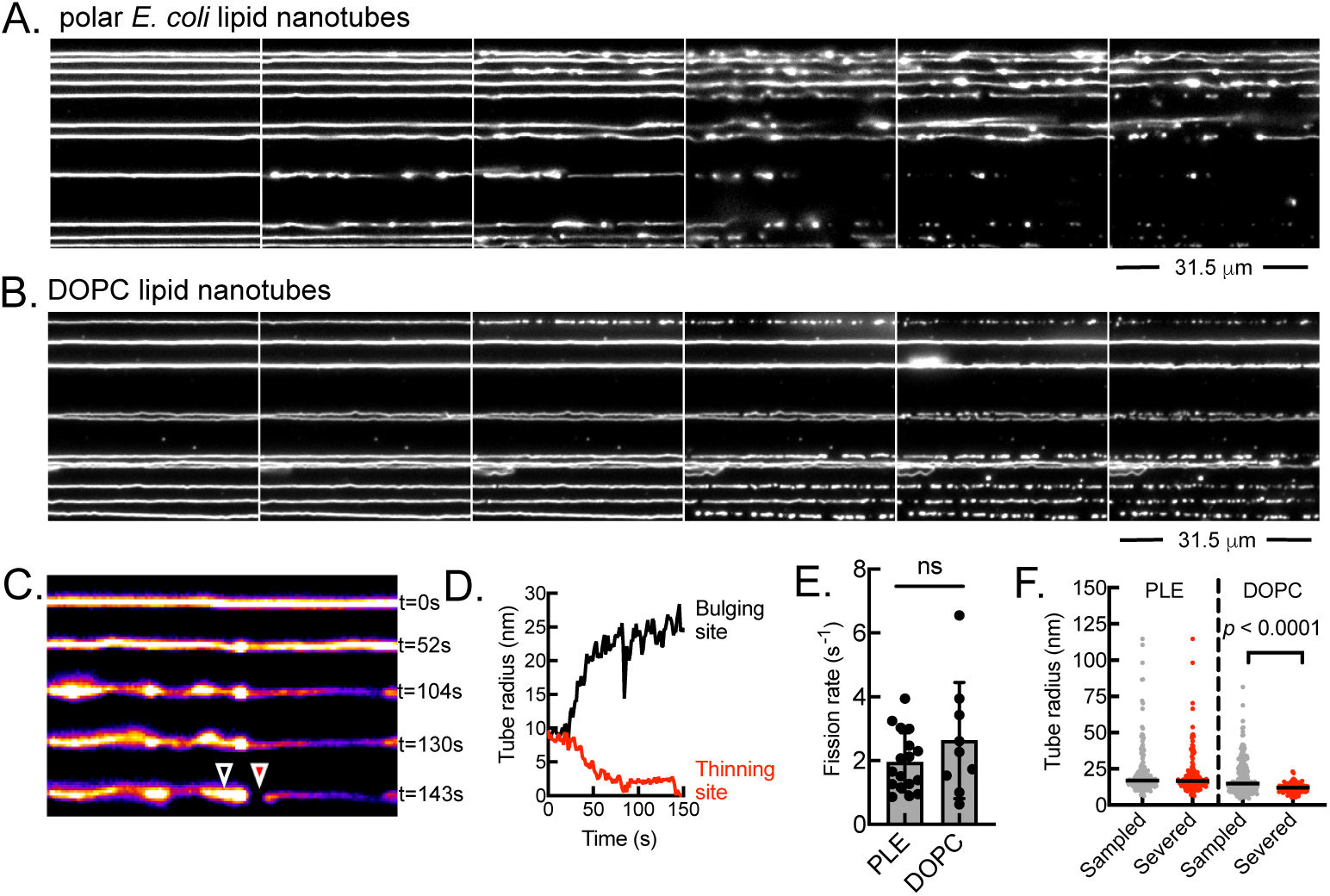
- LL-37 induces fission in A) polar *E. coli* lipid derived supported membrane nanotubes and B) synthetic phosphatidyl choline derived nanotubes. C and D) LL-37 causes transient bulging and thinning of lipid nanotubes concomitant with membrane fission events. E) Polar lipid and POPC-derived nanotubes are disrupted at similar rates F) the presence of polar lipids in nanotube preparations increases the range of nanotube sizes that are disrupted by exposure to LL-37. Fission rates were compared by Student’s t-test while comparisons of nanotube sizes formed to nanotube sizes that underwent fission were compared by Kruskall-Wallis ANOVA with Dunn’s Multiple Comparison post-hoc tests to compare specific groups.

### SMrTs composed of neutral lipids show reduced LL-37-mediated fission

To evaluate the contribution of polar lipids to observed fission activity, we tested LL-37 on neutral DOPC membrane templates. Flowing 50 nM of LL-37 showed no effect on planar bilayers, consistent with the results shown above (Supplemental Fig. S2B), while it caused severing of membrane tubes (Fig 4B). Membrane tubes in these preparations were estimated to be ∼9 ± 4 nm radius (mean ± SD, n = 11, Fig. 4B), similar to that seen with templates prepared from polar lipid extracts (Fig. 4C). While the bulk of the membrane on SMrTs rests on a passive PEG cushion, templates remain tethered to the surface via sparse pinning sites on glass that resist passivation by PEG. This allows monitoring multiple fission events on a single tube, as can be seen in a series of images generated from a time lapse sequence of a single tube exposed to LL-37 (Fig. 4A and 4B). Estimating the cumulative number of tube fragments generated with time and a segmental linear regression analysis of this data provides an estimate of fission kinetics (Fig. 4D, red trace). This analysis across multiple tubes and multiple fission sites reveals similar fission kinetics for PLE and DOPC nanotubes, indicating that LL-37 is equally efficient at severing membrane tubes formed of PLE or DOPC (Fig. 4E), however, comparison of tube diameters created vs the diameter of tubes that undergo fission revealed that for tubes created from PLE, the widest tube severed was ∼115 nm while that with DOPC was only ∼20 nm, demonstrating that polar lipids enhance the fission activity of LL-37 (Fig. 4F).

### The PhoPQ regulated protein, QueE, mediates filamentation and LL-37 resistance

The observation that actively dividing cells are sensitive to LL-37 raised the question of whether stalling division might protect bacteria from killing by this cationic antimicrobial peptide. QueE is a moonlighting protein normally involved in the biosynthesis of the modified base queuosine, but it also stalls *E. coli* division in a PhoPQ-dependent manner (14, 15). We compared the survival of *E. coli* BW25113 and BW25113Δ*queE* to LL-37, the murine orthologue of LL-37, mCRAMP and polymyxin B, a bacterial-derived antimicrobial peptide. We observed reduced survival of the Δ*queE* strain after 10 minutes exposure to both LL-37 and mCRAMP, but not to polymyxin B (Fig. 5A). Consistent with previous results (14, 15), we demonstrated that *E. coli* BW25113 shows *queE* dependent increases in cell length under limiting Mg^2+^ conditions (Fig. 5B). Growth under PhoPQ-inducing conditions also caused bacterial cells from three UPEC strains, SMH38B, CFT073 and UTI89 to exhibit an increased mean cell length (Supplemental Fig. S2). However, growth in low magnesium, did not lead to increased cell length in K12Δ*queE*, confirming that the filamentation we see under PhoPQ-activating conditions is QueE mediated (Fig. 5B). Finally, as previous reports suggested that synthetically filamentous cells are abrogated in colony formation on solid medium (16, 17), we conducted live-cell LL-37 killing assays on *queE*-expressing cells. Briefly, *E. coli* K12, K12*11queE* and K12Δ*queE* + pRL03 bacteria were placed on LB-agarose pads in the presence of propidium iodide before exposing bacteria to LL-37 in water and monitoring bacterial killing, as described above for actively dividing cells. As seen in Fig 5C, we observed significantly enhanced killing of K12Δ*queE* strains in the presence of LL-37 as well as enhanced protection of K1211*queE* expressing QueE from pRL03 cells during exposure to LL-37.

**Figure 5.**
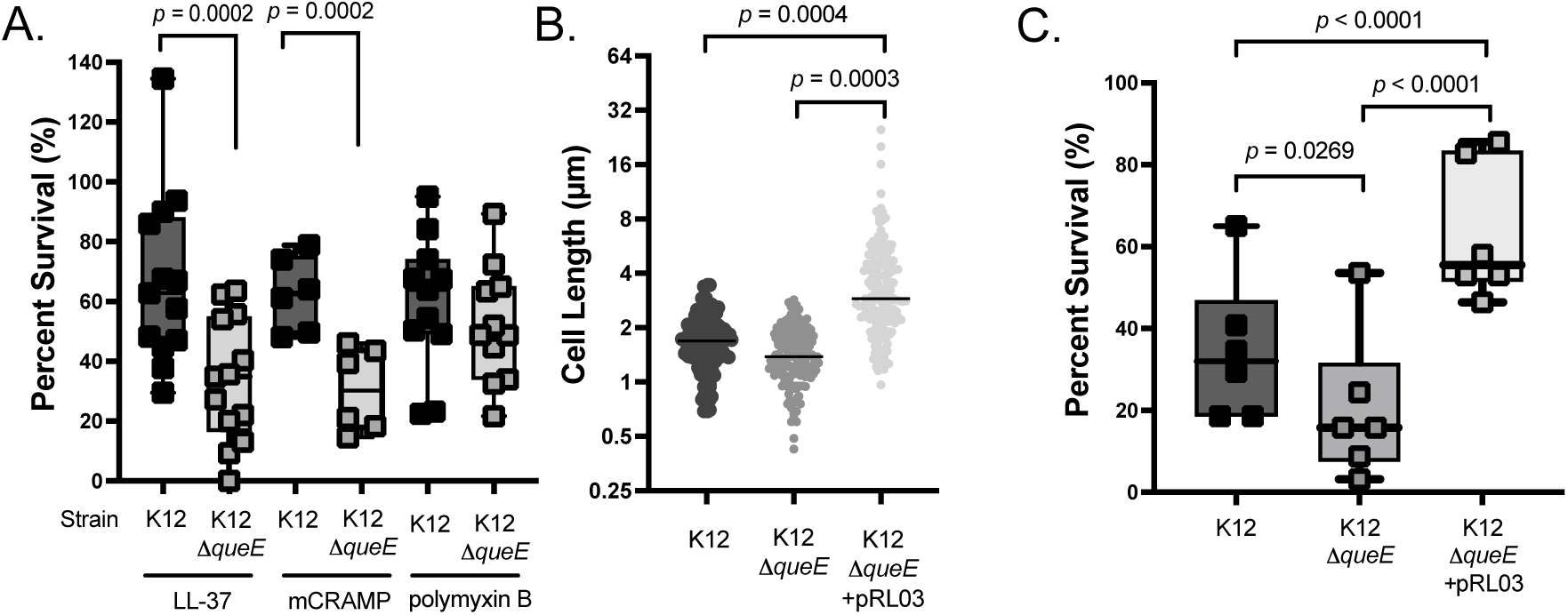
- Deletion of *queE* increases *E. coli* K12 susceptibility to LL-37 and mCRAMP while overexpression of *queE* leads to protection from these molecules. A) Deletion of *queE* results in increased bulk susceptibility to LL-37 and mCRAMP but not to polymyxin B B) Deletion of *queE* leads to reduced *E. coli* K12 mean cell length C) live single-cell killing analysis of *queE*-induced filamentation and killing in *E. coli* K12 shows that cells that are made filamentous via *queE* expression from pRL03 are significantly protected from LL-37 mediated killing. Statistical differences in bulk susceptibility to LL-37, mCRAMP or polymyxin B were assessed by paired t-test of the percent survival following exposure to each compound as assessed by plate counting. Differences in cell lengths were assessed by one-way ANOVA comparing the average cell lengths from three independent experiments (N=50 cells counted for each group). Differences in single cell survival were assessed by comparing the percentage of propidium iodide negative cells from each group following exposure to 100 μg/ml LL-37 by one-way ANOVA with Holm-Šídák’s multiple comparison post-hoc testing.

### Filamentous cells are killed by LL-37 if division resumes

The mechanism of *queE*-mediated filamentation is not well-established but is thought to involve inhibition of divisome maturation (14). As described by Yadavalli et al, *queE*-induced filaments show substantial length heterogeneity, with some cells exhibiting near-normal cell lengths while other cells show dramatic length increases (Supplemental Movie S5) (14). Exposing these populations to LL-37, we observed that killing was observed within the shorter, actively dividing population, while filamentous cell were protected from LL-37 killing, consistent with results shown above Fig 6AB. To determine whether there was a difference in divisome lifespan in QueE-expressing cells, we imaged cells expressing FtsZ-GFP that were also expressing QueE from a compatible, arabinose-inducible plasmid. In general, Z-rings were stable over extended periods of time in the QueE-expressing filamentous cells, showing significantly increased duration compared to those in WT *E. coli* (Fig 6C). However, we observed some uncommon examples where previously stable FtsZ-rings appeared to progress/dim in cells exposed to LL-37, consistent with these cells resuming division. In these cells, the disassembly of FtsZ-GFP was immediately following by propidium iodide entry at the site of Z-ring resolution, similarly to that observed in normally dividing *E. coli* cells (Fig. 6D, Supplemental Movie S6). This is consistent with a model where filamentation no longer protects from killing if divisome function is restored.

**Figure 6.**
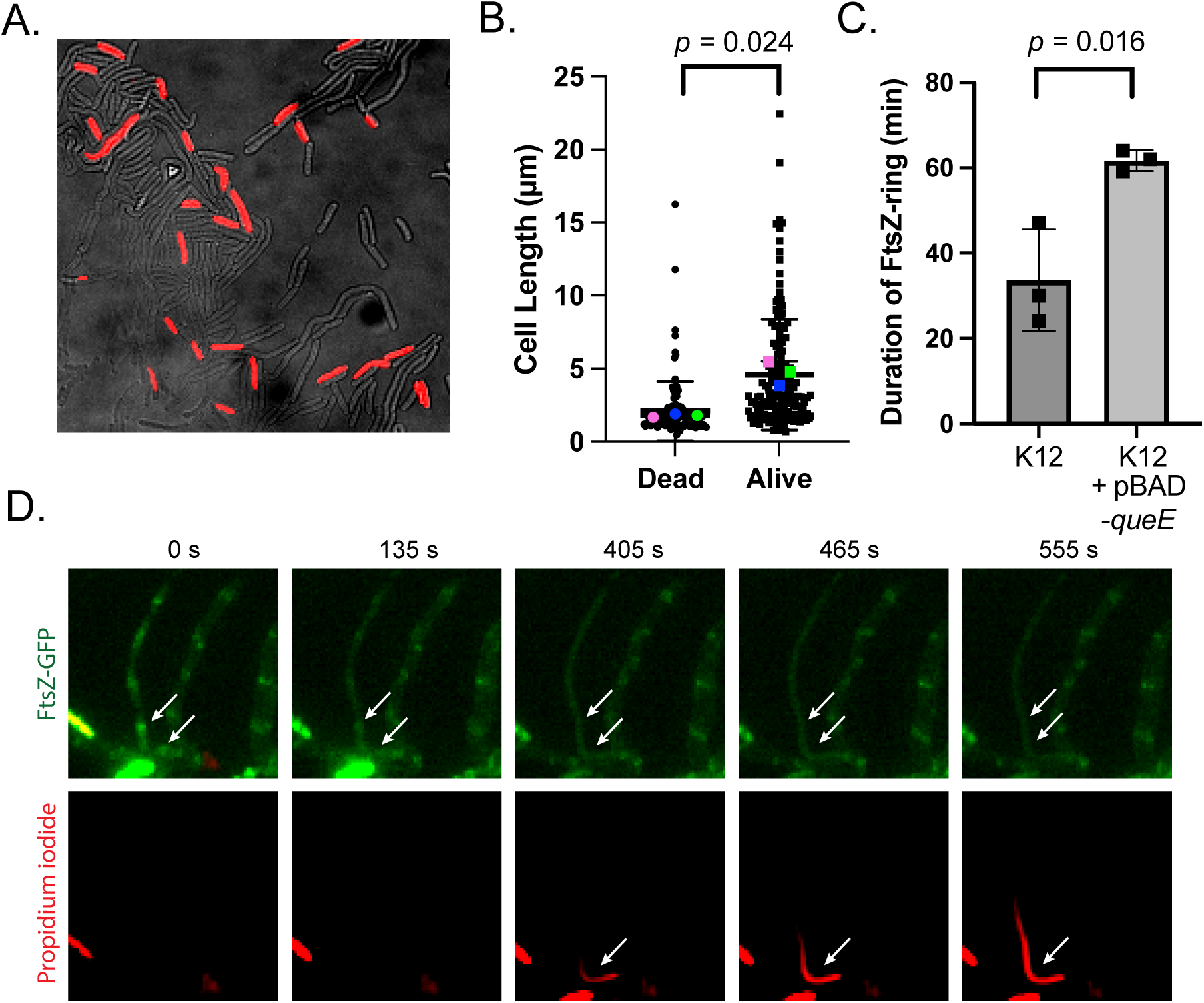
- Killing behaviour of LL-37 in heterogenous populations of *E. coli* K12. A) Induction of filamention from pBAD-*queE* is heterogenous and PI-staining is preferentially observed in the shorter, dividing subpopulation of bacteria B) Cell length analysis of PI-stained LL-37-killed bacteria in K12 + pBAD-*queE* C) Z-ring lifetime analysis in cells expressing QueE shows increased stability of Z-rings D) filamentous cells may be killed by LL-37 whe Z-ring disassembly proceeds (white arrows). Differences in cell length and FtsZ lifetime were assessed by Student’s t-test.

### QueE, SulA and DamX-induced filamentation protect UPEC from LL-37 mediated killing

To determine whether the observations made above were confined to K12 *E. coli* or to QueE-mediated filamentation, we assessed the behaviour of filamentous UPEC strain UTI89 (18) induced via expression of QueE (14, 15), SulA (19) or DamX (20). As shown in Fig 7A, expression of *queE* via growth in limiting Mg^2+^ resulted in a ∼35% increase in mean cell length and this increased depended entirely on functional *queE*. Adding pRL03 to the UTI8911*queE* strain complemented the cell length phenotype, resulting in a 66% increase in mean cell length in low Mg^2+^-grown cells. Importantly, we noted significant protection of the UTI8911*queE* strain when pRL03 was expressed (Fig. 7D). Consistent with previous results, overexpression of either *sulA* (19) or *damX* (20) led to significantly enhanced cell lengths, with a 235% increase when SulA was overexpressed (Fig. 7B), or a 191% increase when DamX was overexpressed (Fig 7C). We then exposed these strains to LL-37 and observed that under conditions where filamentation was observed, we noted significant protection from LL-37 mediated killing (Fig 7D-F).

**Figure 7.**
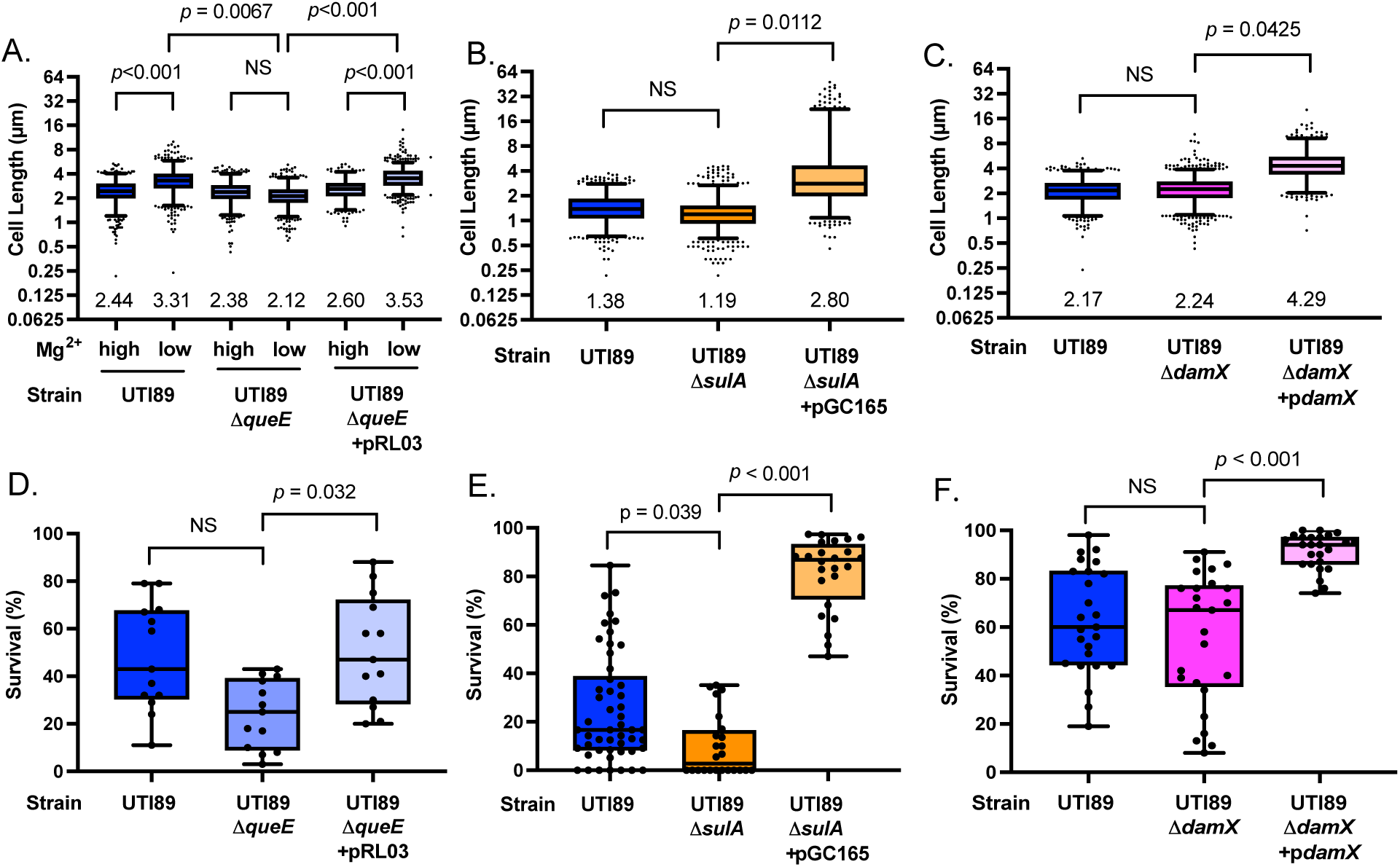
- Filamentation induced by overexpression of QueE, SulA or DamX leads to enhanced LL-37 resistance in uropathogenic *E. coli*. A) QueE mediates low Mg^2+^ induced increased cell length in UTI89 B) Overexpression of the cell division inhibitor SulA increases mean UPEC cell length C) Overexpression of DamX increases mean UPEC cell length. Statistical differences in cell length were assessed by Kruskal-Wallis ANOVA with Dunn’s post-hoc test. The median cell length is indicated on the axis for each strain and condition D - F) filamentation induced by QueE, SulA or DamX increases resistance to LL-37 mediated killing, as assessed by measuring propidium iodide uptake. Statistical significance was assessed by Kruskal-Wallis ANOVA with Dunn’s post-hoc test.

## Discussion

Bacterial killing by the antimicrobial peptide LL-37 is well-studied, and yet the specific mechanism of action remains somewhat enigmatic (21, 22). The peptide is randomly structured in pure water, but forms a kinked amphipathic α-helix in the presence of low concentrations of salt or in trifluoroethanol or dodecylphosphatidylcholine (DOPC) micelles (42, 43). LL-37 associates with bacterial lipopolysaccharide, with the cytoplasmic membrane, and with other intracellular targets, as well as with itself, forming higher-order oligomeric complexes that have been proposed to play a role in antimicrobial activity (23, 24). LL-37 initially interacts with membranes by binding in a parallel orientation to the plane of the membrane, with the aromatic phenylalanines shallowly penetrating the membrane bilayer (7, 25). The nature of the fatty acyl chains also seem to alter peptide/membrane interactions, particularly at very high peptide concentrations (26, 27). This has led to the hypothesis that LL-37 disrupts bacterial membranes via a “carpet” mechanism, whereby peptide-lipid aggregates disrupt the planar membrane bilayer leading to killing. However other model membrane studies have supported both the formation of transmembrane pores (28) or toroidal pores (29, 30), further demonstrating the lack of clarity on how this important molecule helps protect the host.

In the majority of previous studies, the mechanism of LL-37 mediated killing is inferred from studies in static, model membrane systems composed of simplified lipid mixtures. Although valuable, these approaches do not give information about which of the many mechanisms of peptide/lipid interaction actually lead to death in actively growing bacterial cells nor do they monitor behaviour in the more complex lipid environments that typify living bacterial cells. Using live cell imaging, we show that bacterial death, as assessed by Sytox Green or propidium iodide (PI) entry, is not random and clearly occurs in dividing cells downstream of the initiation of septal invagination (Fig 1), consistent with previous results from the Weisshaar group (31). In normally dividing *E. coli*, PI entry occurs at the midline of dividing bacteria and both daughter cells are killed at the same time. As divisome kinetics in *E. coli* are well-characterized, we measured killing as a function of the timing of specific divisome-associated proteins assembly and disassembly from the divisome. FtsZ is the initiator of divisome assembly, forming Z-rings on the cytoplasmic face of the membrane at the site of division anchored to the membrane by FtsA and ZipA. This early assembly is followed, in order, by FtsEX, FtsK, FtsQLB and FtsWI. Finally, FtsN binds to and activates FtsWI activity, leading to activation of the periplasmic divisome complex, septal peptidoglycan biosynthesis and committment to division (32). Thus, FtsN recruitment and activation is a key decision point in cell division.

As division progresses, the FtsZ and FtsN rings constrict (33), reducing the diameter of both as an inwardly oriented wedge of septal PG is synthesized (34). This continued inward PG synthesis eventually becomes a major force generation factor in the progression of the divisome itself and the FtsZ ring is disassembled before cytoplasmic membrane scission is completed (33–35). FtsN disassembly follows shortly thereafter in two distinct stages, although it is unclear whether inner membrane scission is complete at the time of FtsN disassembly (33–35). In our results, we noted that FtsZ and FtsN were almost completely absent at the time of PI entry to LL-37 exposed bacterial cells (Fig. 2A-D), suggesting that LL-37 mediated killing occurs in very late bacterial division.

In parallel with sPG synthesis, the amidases, AmiA, AmiB and AmiC, are recruited to the divisome, where they contribute to the final cleavage of septal PG, thereby allowing daughter cell separation. Amidase activity is tightly controlled by activator proteins, ensuring that sPG assembly precedes hydrolysis (36). The amidases have some redundancy, both in terms of activation and localization, but AmiB recruitment to the divisome depends on prior FtsN recruitment and AmiB appears to arrive at the divisome later than AmiA or AmiC (36, 37). Analysis of the assembly dynamics of a catalytically inactive, but appropriately localized, version of AmiB (37), showed that while AmiB was typically disassembled from the divisome at the time of killing, there is a proportion of PI+ bacteria in which an AmiB-sfGFP ring is observed. This is consistent with a model where LL-37 induced death follows membrane constriction due to divisome activity (Fig. 1), downstream of FtsZ and FtsN assembly and dissasembly (Fig. 3, Supplemental Movies S2-S4), consistent with LL-37 targeting actively dividing bacteria for killing.

Introducing polarly localized division sites, either through overexpression of FtsZ or through deletion of the *minCD* system, led to LL-37 mediated membrane permeabilization via polar divisomes. This only occured if the divisomes were productive, as we observed no killing at Z-ring localized sites, unless the division site progressed through later divison events involving FtsZ and FtsN dissembly. Filaments with visible Z-rings could survive in the presence of LL-37 for >10 minutes, as long as that site did not advance beyond FtsZ assembly. However, if the Z-ring progressed, even previously LL-37 resistant filaments could be killed (Fig. 6, Supplemental Movie S4). Thus, division-associated killing is not a consequence of the location of the division site but rather is a consequence of divisome maturation and progression, concomitant with cytoplasmic membrane fission.

The pore-forming activity of LL-37 has been attributed to both electrostatic and hydrophobic interactions between the peptide and membrane lipids but previous reports have described killing occurring via a carpet-like mechanism, a barrel-stave pore model or via toroidal pores (28–30, 38), but more recent reports have described the formation of fibrillar lipid-peptide aggregates that might reflect other modes of action (21, 22, 26). SMrT templates have been used previously to analyze the role of fission proteins in functions of eukaryotic membrane dynamics and in mitochondrial divisome function (13, 39). Here, we used them to show that LL-37 could rapidly induce membrane fission in tubular membrane templates (Fig. 4A-C). Consistent with previous results, we noted that LL-37 interacted with both neutral and charged bilayers (7) but we observed a pronounced diameter dependent relationship to LL-37 induced fission in uncharged DOPC tubes with thinner regions showing increased likelihood of fission while large regions did not undergo LL-37 induced fission (Fig. 4) (25). This relationship between diameter and likelihood of fission disappears when polar *E. coli* lipid extracts are used to create the tubes, suggesting that the presence of charged membrane lipids at the site of fission may increase LL-37 susceptibility (40). Charged lipids are enriched at both bacterial poles as well as at the site of membrane invagination during division, providing plausible localized lipid composition differences that might show altered susceptibility to LL-37 association or perturbation (40, 41). Our results are consistent with previous results from the Weisshaar group showing that dividing cells are actively killed by LL-37 with subsequent alteration of cytosolic diffusion properties (23, 31). This provides a significant advance in the paradigm for how LL-37 interacts with dynamic membrane structures and supports a model whereby division-associated membrane thinning and lipid composition is a key element of LL-37-mediated bacterial killing.

The divisome is critical for septal PG biosynthesis and division however, a number of microbes can stall bacterial division by interfering with normal divisome assembly and progression, either in response to antibiotic treatment, DNA damage or other infection-induced stressors or signals, leading to bacterial filamentation (42–48). Several protein inducers of filamentation have been previously reported in *E. coli*, including one that is transcriptionally activated by antimicrobial peptides themselves. Exposure to the synthetic antimicrobial peptide C18G signals via the PhoPQ two-component regulatory system to induce *queE* mediated division block (14, 15). Divisome-regulated peptidoglycan biosynthesis is a common antibiotic target, highlighting the importance of bacterial control over the timing of division and numerous observations suggest that division blockage is observed during certain types of infection, including *Salmonella enterica* intramacrophage growth, and during urinary tract infection in both *E. coli* and in *Klebsiella pneumoniae* (49–51). The observation that division itself sensitizes bacteria to LL-37 mediated killing suggested that division blockage might also affect susceptibility to LL-37-mediated killing.

During UTI, UPEC initially attach to the surface of bladder epithelial cell via the mannose-sensitive FimH adhesin (52). Binding leads to endocytic uptake by bladder epithelial cells before vaculolar escape leading to cytoplasmic invasion (53). UPEC then proliferate intracellularly to form intracellular bacterial communities (IBCs) creating biofilm-like colonies (7, 8). Within these IBCs, a subpopulation of UPEC adopt a dramatic filamentous morphology coincident with vacuolar breakage and bacterial escape (18). The filamentous and other bacteria escape the intracellular environment and enter the bladder lumen where they can colonize other adjacent sub-populations of bladder epithelial cells (18). UPEC filamentation has been associated with increased infection and bacteria that cannot filament result in reduced infection in murine models of UTI (19).

Urinary tract infections caused by UPEC are among the most common bacterial diseases globally and UPEC have adopted various mechanisms through which they can invade and replicate in bladder epithelial cells to evade or resist host defences (55). During UPEC infection, a number of components of the innate immune system become activated due to the secretion of cytokines and peptides from the epithelial cell (56). Recruitment of mast cells, macrophages and neutrophils to the site of infection leads to localized inflammation and secretion of antimicrobial peptides both from the inflamed epithelium and from the recruited phagocytes (57–59). UPEC strains appear to have specific adaptations to modulate innate immune signaling and to resist killing by these recruited peptides and cells (60–64). UTI leads to increased production of LL-37 in by the bladder epithelium (65, 66). UPEC have specific adaptations to protect them from the antimicrobial activity of LL-37 and other antimicrobial peptides. UPEC strains have specific resistance mechanisms to these innate defences, including production of curli, production of outer membrane proteases, and enhanced biofilm formation (63, 66–68).

The specific signals and signaling systems that drive UPEC filamentation in vivo are not well-characterized, but UPEC filamentation can be induced *in vivo* via the overexpression of the SOS-response effector, *sulA* (19, 69). The SOS-response is activated during bacterial starvation or antibiotic-induced DNA damage leading to a halt in cell division to promote DNA repair mechanisms (70). During DNA damage, RecA is activated and cleaves repressor LexA resulting in the transcription of SOS genes including *sulA,* which directly inhibits FtsZ polymerization (71, 72). Recent work suggests that blocking the SOS-response in UPEC might be a viable path to blocking division stalling, but there is some controversy here as other work suggests that SulA is not a major driver of filamentation during normal in vivo growth (20). Further research is needed to validate these hypotheses in vivo (73).

DamX is an inducer of filamentation in UPEC (20). While the exact mechanism of *damX* mediated filamentation remains unclear, it is recruited to the site of division in an FtsZ dependant manner, consistent with it being a SPOR domain protein that is presumably recruited to sites of septal PG biosynthesis (74). During UPEC filamentation, DamX is uniformly localized across the membrane of filaments and becomes more concentrated at future sites of division, potentially assisting in resumption of cell division of filamentous cells (75). Whether DamX division associated relocalization is a cause or a consequence of division competence remains unclear, but our results demonstrate clearly that DamX-induced filaments are protected from LL-37 killing (Fig. 4).

Growth under PhoPQ-inducing conditions increases cell length in non-pathogenic K12 as well as in three different UPEC strains, (Fig. 5, Supplemental Fig. S2). QueE-mediated CAP resistance is not specific to LL-37, as we also noted altered susceptibility to the murine orthologue of LL-37, mCRAMP (but not to polymyxin B). Work in non-pathogenic *E. coli*, and our results here, have shown that exposure to LL-37 and/or signals that activate PhoPQ, can drive a QueE-dependent stalling of bacterial division (14, 15). Regardless of the specific mechanism by which division stalling/filamentation occurs, filamentation provides UPEC with enhanced protection against LL-37 mediated killing. Filament protection is not due to an inherent imperviousness to LL-37, but rather due to stalled divisome maturation, as filamentous cells induced by QueE overexpression can be killed, if divisome maturation proceeds (Supplemental Movie S6) and minicell producing filaments are also killed from the polarly localized divisome (Fig. 3 and Supplemental Movie S3).

In spite of the apparent importance of filamentation in the UPEC lifestyle during UTI, little is known about the biological consequences of bacterial filamentation or how it contributes to pathogenesis. Filamentation protects numerous types of bacteria from uptake and killing by phagocytic cells, suggesting that enhanced resistance to innate immune responses might be a consequence of bacterial filamentation (19, 76). Dividing *E. coli* are more sensitive to LL-37 mediated killing and during UTI, the antimicrobial peptide LL-37 is highly elevated, serving as a protective factor against continued bacterial invasion (31). Previous work has also demonstrated that there are stochastic differences in LL-37 binding to individual cells within a bacterial population, such that cells that randomly bind to larger numbers of peptide molecules are killed first and that this heterogenous peptide absorption leads to increased survival of the remaining cells that bound less peptide (77). Our results suggest that bacterial division stalling during UTI may represent a protection mechanism from innate-immune related LL-37.

In addition to genetic and environmental factors, some antibiotic treatments can also stall bacterial division, and this has been observed during UTI suggesting that this treatment might, somewhat paradoxically, reduce the effectiveness of host innate immune responses (62). UTIs are typically treated with antibiotics, but increased levels of antibiotic resistance have led to increased rates of treatment failure, highlighting the need for alternate therapeutics against this common type of infection (78). The observed change in morphology to a filamentous form that is associated with escape from IBCs may protect UPEC from components of both cellular and humoral innate immunity. This further suggests that identifying small molecules that can block infection-related filamentation might provide a viable path for potentiating innate immune responses to UTI and that this may represent a novel path to treatment for UTI that could have significant impact on patient outcomes.

## Materials and Methods

### Bacterial Strains and Culture Conditions

The bacterial strains and plasmids used in this study are described in **Table S1.**

### LL-37 Killing Assays

Bacterial cultures were grown overnight in lysogeny broth or N-minimal media containing either high (10 mM) or low (10 µM) MgSO_4_ concentrations. Bacteria were then grown to late-log (OD_600_ ∼ 0.8) and resuspended in 10 mM HEPES buffer, pH 7.5. Cultures were diluted to a final a concentration of 10^7^ cells/ml. Cultures were treated with 50 µg/ml LL-37 for 1 hour and resuspended in PBS. For bulk killing assays, surviving bacteria were plated on LB agar plates while for live-cell killing assays, cells were stained with 10 µg/ml Sytox Green or 1 µg/ml propidium iodide prior to live-cell imaging.

### Confocal Fluorescence Microscopy

Live cell microscopy images were acquired using three different microscopes. The Quorum Spinning disc confocal microscope based on the Quorum DisKovery Multimodal imaging system using the iXon 897 EMCCD camera from Andor and the Metamorph program was used to quantify filamentation and LL-37 survival in *E. coli*. The Olympus Deconvolution microscope using the DP72 camera and based on the CellSens program was also used to quantify filamentation and LL-37 survival in UPEC. The ReScan Confocal microscope based on the Volocity acquisition software was used to capture live-cell timelapse videos of LL-37 killing. The 63x oil-immersion lens was used for all microscopes. DAPI (405nm), green (488 nm), red (561 nm) and far red (637 nm) lasers were all used with appropriate fluorophores. For live-cell division and LL-37 killing timelapse videos, a heated stage was used to maintain a constant temperature (37°C) of the chamber. The CO_2_ concentrations were kept at zero for all experiments.

### Supported Membrane Templates (SMrTs)

Total lipid concentration of the *E. coli* Polar Lipid Extract (PLE) (Avanti Polar Lipids) was calculated based on the reported distribution of 67% (w/w) phosphatidylethanolamine, 23.2% (w/w) phosphatidylglycerol and 9.8% (w/w) Cardiolipin on the company website. SMrTs were prepared as described earlier (31). Briefly, lipid mixes of PLE and Texas Red DHPE (Invitrogen) were reconstituted in a 99:1 molar ratio in chloroform and brought to a final concentration of 1 mM total lipid. The same was done for 1,2-dioleoyl-glycero-sn-3-phosphocholine (DOPC, Avanti Polar Lipids). 2 μL of the lipid mix was spread with a glass syringe on a PEGylated glass coverslip, dried and assembled inside an FCS2 flow chamber (Bioptechs). Lipids were hydrated at room temperature by filling the chamber with 20 mM HEPES pH 7.4 buffer with 150 mM NaCl. SMrTs were prepared by flowing buffer over the dry lipid mix, which produced planar lipid bilayer islands where the lipid mix was spotted and an array of membranes tubes downstream. Stock solution of 1 mM LL-37 (Catalog. no HY-P1222, MedChemExpress) was prepared in deionized water and stored at −80°C. The LL-37 stock was diluted to 50 nM in 20 mM HEPES pH 7.4 buffer with 150 mM NaCl and used for all experiments. Templates were imaged through a 100x, 1.4 NA oil-immersion objective on an Olympus IX83 inverted microscope connected to an LED light source (CoolLED) and an Evolve 512 EMCCD camera (Photometrics). Image acquisition was controlled by µManager and images were analyzed using Fiji. Tube sizes were estimated based on a calibration procedure described earlier (31). Movies of LL-37 flown on to membrane tubes were acquired at 132 ms time interval. ROIs marking single tubes were isolated from these movies. The number of tube fragments at every frame was estimated from a particle analysis routine that involved creating a mask using the default threshold setting. Segmental linear regression analysis (Graphpad Prism) of the cumulative particle number plotted against time provided estimates of fission rate.

### Statistical Analysis

All experiments were performed at least three independent times with the specific total number indicated in the respective figure legends along with sample sizes. Statistical analysis performed using GraphPad Prism 10 for MacOS with the actual tests disclosed in the respective figure legend. Non-parametric tests were chosen for comparisons of cell length as data from these experiments were typically not normally distributed. For comparisons of cell survival, normality was assumed and data was typically analyzed with paired Student’s t-test when comparing two populations, with a repeated-measures one-way ANOVA for three or more samples with one parameter, or repeated-measures two-way ANOVA when two parameters were considered. Post-hoc tests were employed based on assumptions and comparisons as recommended by GraphPad Prism and disclosed in the respective figure legends when comparing multiple conditions. Here, we accept *p*<0.05 as an indication of statistical significance, but we report *p* values for greater transparency.

## Acknowledgements

The Natural Sciences and Engineering Research Council funded this research (RGPIN-2024-05155) with contributions from Toronto Metropolitan University Faculty of Science (36803) and Office of the Vice-President of Research and Innovation (36812) with awards to JBM. The Canadian Institutes of Health Research funded this research with awards to RJB (PJT-183914), the Canada Fountain for Innovation (32957, 38151), and the Canada Research Chairs Program (950–232333) with contributions from Toronto Metropolitan University (38151) to RJB.

---

**Table S1.**
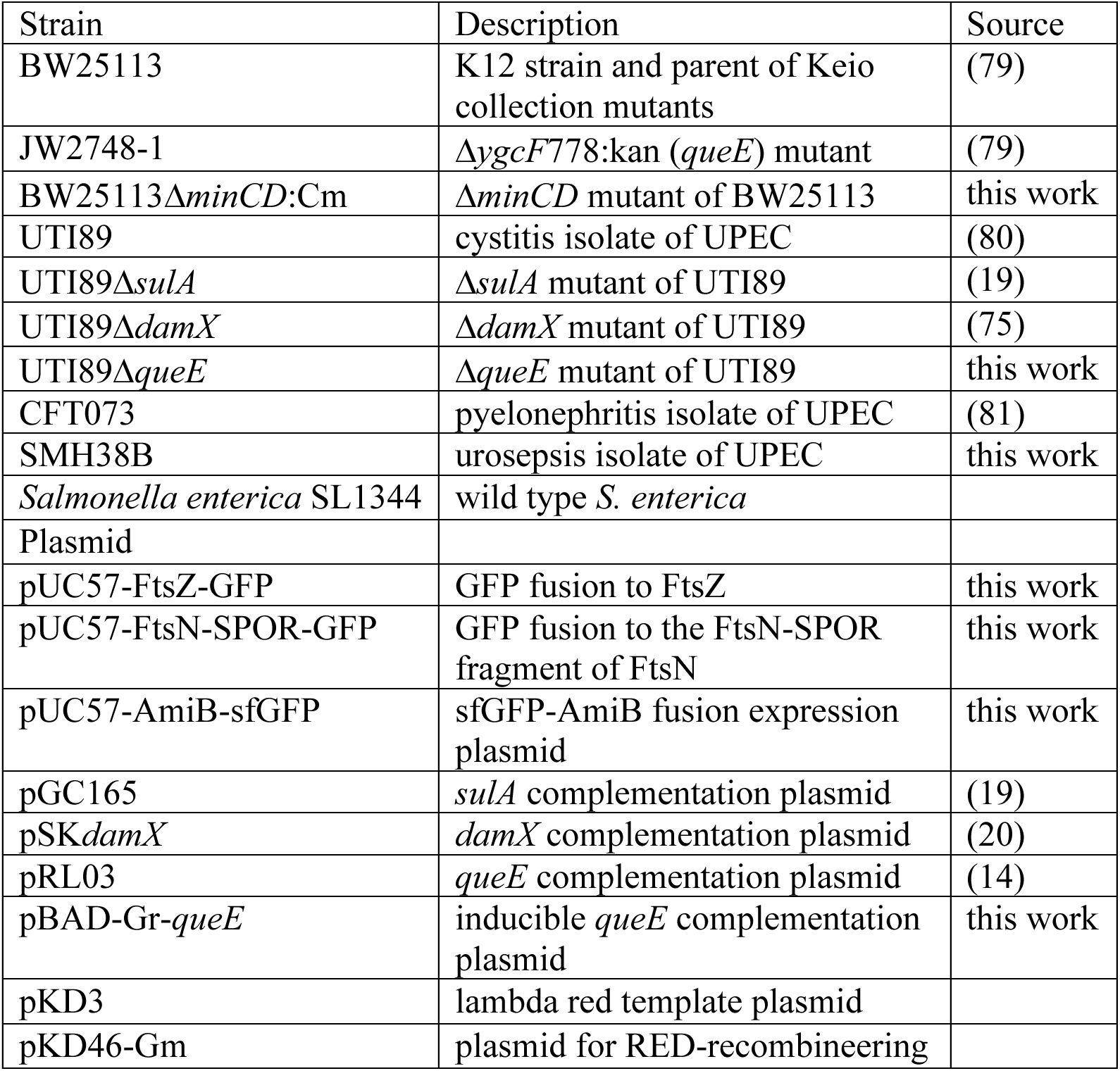
Strains and plasmids used in this study.

