## Supplemental Information for "Division-arrest induced filamentation protects uropathogenic *Escherichia coli* from killing by the cathelicidin antimicrobial peptide LL-37"

### SI Materials and Methods

#### Bacterial Strains and Culture Conditions

Cultures were grown routinely in lysogeny broth (LB) or N-minimal media containing 0.1% casamino acids, 0.2% glucose, 0.002% thiamine, 15 μM iron sulfate, supplemented with either high (10 mM) or low (10 μM) $MgSO_4$ concentrations. Cultures were grown at 37°C with shaking at 200 rpm. Antibiotics for selection were used at the following concentrations: ampicillin 100 μg/ml, kanamycin 50 μg/ml and chloramphenicol 25 μg/ml. The *lac* and *trc* promoters were induced with 0.5 mM β-isopropyl-D-thiogalactoside (IPTG) and the *araBAD* promoter was induced with 0.2% arabinose.

#### Molecular cloning

Creation of UTI89Δ*queE*-kanR: A *queE*::kanR allele flanked upstream and downstream by the upstream and downstream regions of the *queE* gene in *E. coli* UTI89 was synthesized by GenScript. This cassette was then subcloned to pRE112-Gm as a SacI-KpnI fragment. The pRE112-*queE*-Kan plasmid was mobilized to competent UTI89 + pKD46 via conjugation from *E. coli* S17-1λ*pir*, recovered for 12 hours at 30°C and then plated on LB containing 100 μg/ml ampicillin and 50 μg/ml kanamycin. Merodiploid strains were counterselected on sucrose and gentamicin-sensitive; kanamycin-resistant clones were identified by selective plating.

BW25113Δ*minCD* was created via the lambda-red deletion system of Datsenko and Wanner. Putative ΔminCD strains were screened by colony PCR before being assessed for altered morphology and behaviour.

For the expression of fluorescently labelled fusion proteins, briefly, FtsZ-GFP, FtsN-GFP or GFP-AmiB fusions were designed based on published studies and synthesized by GenScript. All fusions were expressed from pUC57-Amp and these showed basal level expression that did not interfere with observable normal bacterial physiology.

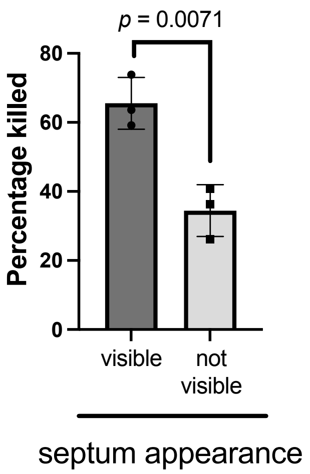


Supplemental Figure S1. Dividing *Salmonella enterica* SL1344 are specifically sensitized to killing by LL-37. The percentage of cells with visible vs non-visible septa were assesed for death by propidium iodide uptake following exposure to 10 μg/ml LL-37. Significance determined by Student's t-test, N = 249 to 1370 cells counted per experimental replicate.

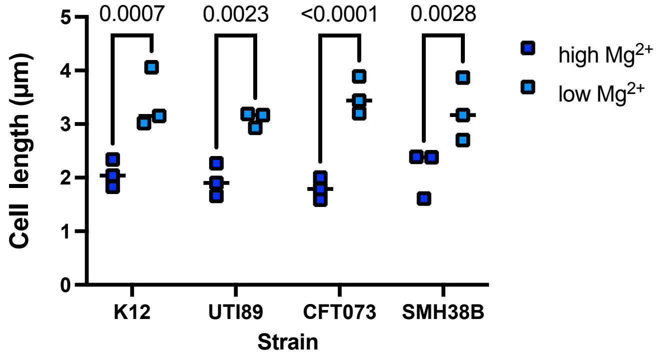


Supplemental Figure S2. Growth under PhoPQ-inducing conditions caused increased mean cell length in both K12 strains and in three UPEC strains - SMH38B, CFT073 and UTI89. At least 50 cells were measured for each replicate and statistical significance was determined by student's t-test.

| Strain | Description | Reference |
|---|---|---|
| *E. coli* BW25113 | WT K12 *E. coli* | (79) |
| E. coli JW2748-1 | K12Δ*queE*:KanR | (79) |
| E. coli JW1116-1 | K12Δ*phoP*:KanR | (79) |
| BW25113 Δ*queE* | K12Δ*queE* | this publication |
| BW25113 Δ*phoP* | K12Δ*phoP* | this publication |
| UTI89 | UPEC cystitis isolate | (80) |
| UTI89Δ*queE* | deletion of *queE* in UTI89 strains | this publication |
| UTI89Δ*sulA* | deletion of *sulA* in UTI89 strain | (19) |
| UTI89Δ*damX* | deletion of *damX* in UTI89 strain | (20) |
| CFT073 | UPEC urosepsis isolate | (67) |
| SMH38U | UPEC pyelonephritis/urosepsis isolate | this publication |
| Plasmid | Description | Reference |
| pUC57-Amp | cloning plasmid | Genscript |
| pRL03 | *queE* expressed in pTrc99a | (14) |
| pBADGr-*queE* | *queE* expressed from pBADGr | this publication |
| pGC165 | *sulA* expressed in pUC8 | (19) |
| pSK*damX* | *damX* expressed in pBAD33 | (20) |
| pUC57-FtsZ-GFP | FtsZ-GFP fusion expression plasmid | this publication |
| pUC57-FtsN-SPOR-GFP | FtsN SPOR domain-GFP expression plasmid | this publication |
| pUC57-AmiB-sfGFP | AmiB-sfGFP fusion expression plasmid | this publication |
| pKD3 | lambda red template plasmid | |
| pKD46-Gm | plasmid for RED-recombineering | |

Supplemental Table S1. Strains and plasmids used in this study

Supplemental Movie S1. Dividing BW25113 are killed by LL-37 downstream of FtsZ ring assembly and dissassembly.

Supplemental Movie S2. Dividing BW25113 are killed by LL-37 downstream of FtsN ring assembly and dissassembly

Supplemental Movie S3. Dividing BW25113 are killed by LL-37 downstream of AmiB ring assembly and dissassembly

Supplemental Movie S4. LL-37 can target and disrupt polarly localized divisomes that are observed adjacent to minicells produced in BW25113Δ*minCD* strains.

Supplemental Movie S5. Effect of LL-37 on *E. coli* polar lipid extract derived SMrTs.

Supplemental Movie S6. Preferential killing of dividing/shorter subpopulation within QueE-overexpressing bacteria. Heterogenous-sized populations of *E. coli* BW25113 were created by expressing QueE from the *araBAD* promoter in pBAD-Gr-*queE*. Cells were exposed to LL-37 as described in the materials and methods.

Supplemental Movie S7. Close up of LL-37 mediated killing in QueE-induced filamentous cells that also contained pUC57-FtsZ-GFP. We noted sensitization of QueE-induced filamentous *E. coli* cells to LL-37 when divisome progression resumes.
